# The expansion in lymphoid organs of IL-4^+^ BATF+ T follicular helper cells is linked to IgG4 class switching *in vivo*

**DOI:** 10.1101/284737

**Authors:** Takashi Maehara, Hamid Mattoo, Vinay S. Mahajan, Samuel J.H. Murphy, Grace J. Yuen, Noriko Ishiguro, Miho Ohta, Masafumi Moriyama, Takako Saeki, Hidetaka Yamamoto, Masaki Yamauchi, Joe Daccache, Tamotsu Kiyoshima, Seiji Nakamura, John H. Stone, Shiv Pillai

## Abstract

Distinct T_FH_ subsets that influence specific class-switching events are assumed to exist, but the accumulation of isotype-specific T_FH_ subsets in secondary and tertiary lymphoid organs has not been hitherto demonstrated. IL-4 expressing T_FH_ cells are surprisingly sparse in human secondary lymphoid organs. In sharp contrast, in IgG4-related disease (IgG4-RD), a disorder characterized by polarized Ig class switching, most T_FH_ cells in tertiary and secondary lymphoid organs make IL-4. Human IL-4^+^ T_FH_ cells do not express GATA-3 but express nuclear BATF, and the transcriptomes of IL-4 secreting T_FH_ cells differ both from PD1^hi^ T_FH_ cells that do not secrete IL-4 and IL4-secreting non-T_FH_ cells. Unlike IgG4-RD, IL-4^+^ T_FH_ cells are rarely found in tertiary lymphoid organs in Sjögren’s syndrome, a disorder in which IgG4 is not elevated. The proportion of CD4^+^IL-4^+^BATF^+^ T cells as well as of CD4^+^IL-4^+^CXCR5^+^ T cells in IgG4-RD tissues correlates tightly with tissue IgG4 plasma cell numbers and plasma IgG4 levels in patients but not with the total plasma levels of other isotypes. These data describe a disease-related T_FH_ sub-population in human tertiary and secondary lymphoid organs that is linked to IgG4 class switching.

## INTRODUCTION

T follicular helper (T_FH_) cells provide help to B cells during T-dependent immune responses and they contribute to isotype switching, somatic hypermutation, germinal center formation and the selection of high affinity B cells in the germinal center (Crotty, 2011; King et al., 2008; Vinuesa et al., 2005). Exactly how T_FH_ cells provide specificity to class switching events is unclear. The idea that unique T_FH_ subsets separately and specifically drive class switching to different Ig isotypes is attractive, but no *in vitro* or *in vivo* data exist to firmly establish this notion. Indeed, there have been no studies using multicolor staining approaches to examine human T follicular helper cells *in situ* in secondary or tertiary lymphoid organs. The possibility that chronic disease states exhibiting polarized isotype switching could provide novel insights about specialized T_FH_ cells served as the rationale for undertaking this study.

Some evidence for specialized T_FH_ subsets, albeit indirect, comes from studies of circulating human T_FH_ cells that have described three T_FH_ subsets defined on the basis of chemokine receptor expression patterns. The relationship between blood T_FH_ cell subsets and T_FH_ cells in secondary or tertiary lymphoid organs remains unclear. In the studies of Ueno et al. (Morita et al., 2011; Ueno et al., 2015) on blood T_FH_ subsets, T_FH1_ cells secrete IFN-γ upon activation and have limited isotype-switching activity when examined in *in vitro* co-culture experiments. T_FH2_ cells secrete IL-4 after many days of *in vitro* stimulation and can mediate class switching to IgA, IgE and essentially all IgG isotypes, including IgG4. T_FH17_ cells secrete IL-17 following activation and are equally promiscuous. Although all T_FH_ cells may have the potential to secrete IL-4, one report has described polarized IL-4 secreting T_FH_ cells in mice in the context of an allergic disease model, and it was suggested that these cells could subsequently differentiate into T_H2_ cells (Ballesteros-Tato et al., 2016). An illuminating study using reporter mice has led to the view that T_FH_ cells initially make IL-21, mature into cells that make IL-21 and IL-4, and eventually make IL-4 alone (Weinstein et al., 2016). These studies also demonstrated that the use of a Type 2 response-linked murine pathogen facilitated the induction of IL-4 secreting “T_FH4_” cells. There have been no other reports establishing the existence of functionally distinct T_FH_ subsets in human or murine secondary or tertiary lymphoid organs. Moreover, no cytokine-expressing subset of these cells in tissue sites has been linked so far to any specific disease, nor have T_FH_ subsets been defined that determine specific polarized class-switching events. How the overall transcriptome of an IL-4 secreting T_FH_ cell population may differ from other T_FH_ cell types has also never been determined, since such cells have not previously been purified from secondary or tertiary lymphoid organs.

IgG4-related disease is a chronic inflammatory disease characterized by tumescent lesions with characteristic storiform fibrosis, obliterative phlebitis and a marked lymphoplasmacytic infiltrate that includes a large proportion of IgG4-positive plasma cells (Kamisawa et al., 2015; Mahajan et al., 2014). Circulating expansions of plasmablasts, most of which express IgG4, are a hallmark of active disease (Mattoo et al., 2014). We have shown that circulating plasmablasts are heavily somatically hypermutated, implying that these B lineage cells are derived from germinal centers. We have also shown that subjects with IgG4-RD exhibit large clonal expansions of CD4^+^ CTLs, the dominant T cells in disease tissues, and that these cells are activated at lesional sites, where they secrete IL-1β, TGF-β1 and IFN-γ (Mattoo et al., 2016; Maehara et al., 2017). Although an increase in blood T_FH2_ cells has been noted in IgG4-RD (Akiyama et al., 2015; Akiyama et al., 2016), these cells function promiscuously *in vitro*, as they can facilitate class switching to multiple isotypes in co-culture experiments. There is no evidence so far connecting blood T_FH_ cell subsets to any functional counterparts in secondary or tertiary lymphoid organs. Multi-color staining approaches have hitherto not been used to examine lymphoid organ T_FH_ cells *in situ*. We show here, using such an approach, that T_FH_ cells making IL-4 are surprisingly sparse in normal human tonsils and mesenteric lymph nodes. However, IL-4 and BATF expressing T_FH_ cells are dramatically expanded in IgG4-RD tertiary lymphoid organs and lymph nodes, frequently making up more than half of all T_FH_ cells. This subset of human T_FH_ cells can be found in association with AID expressing B cells, is more frequent in extra-follicular sites than in the light zone, and is linked to specific class switching to IgG4.

Isolating T cell subsets based on their function, such as secretion of their cardinal cytokine, and characterizing their gene expression profiles can help us gain better insights into the biology of specific T_FH_ subsets, as well as define better surface markers for their flow cytometric identification. We therefore used an IL4-capture assay followed by FACS sorting, to isolate IL4-secreting T_FH_ cells from a human tonsil and compared their transcriptomic profiles with CXCR5^hi^PD1^hi^IL4^-^ tonsillar T_FH_ cells and IL4-producing CXCR5^-^ non-T_FH_ cells (T_H2_ cells). Our studies validate the notion of functionally distinct T_FH_ subsets, establish a link between tissue and lymphoid organ human IL-4 secreting T_FH_ cells and IgG4-related disease, and identify genes that are specifically expressed in and define the human IL-4 secreting T_FH_ cell subset.

## RESULTS

### CD4^+^ICOS^+^IL-4^+^ T cells are sparse in normal tonsils and lymph nodes

We initially examined normal tonsils to quantitate CD4^+^ICOS^+^ T cells that express IL-4 *in situ*. Almost all the IL-4 expressing cells seen in tonsils were CD4^+^, but did not express ICOS (Fig. 1, A and B). These CD4^+^IL-4^+^ICOS^-^ T cells are presumably T_H2_ cells that reside in T cell zones in tonsils. Quantitation revealed that CD4^+^ICOS^+^IL-4^+^ T_FH_ cells represent less than 0.5% of all CD4^+^ICOS^+^ T_FH_ cells in normal tonsils and lymph nodes (Fig. 1C). On the other hand, quantitative analyses revealed that about 40% of CD4^+^ICOS^+^ T_FH_ cells in an IgG4-RD subject expressed IL-4 (Fig. 1C). The low frequency of T_FH_ cells that express IL-4 was confirmed in twelve different tonsil specimens (Fig. 1D). Relatively low frequencies of IL-4^+^CD4^+^ICOS^+^ and IL-4^+^ICOS^+^Bcl6^+^ T_FH_ cells were also observed in normal mesenteric lymph nodes in addition to tonsillar samples (Figs 1E).

**Fig. 1.**
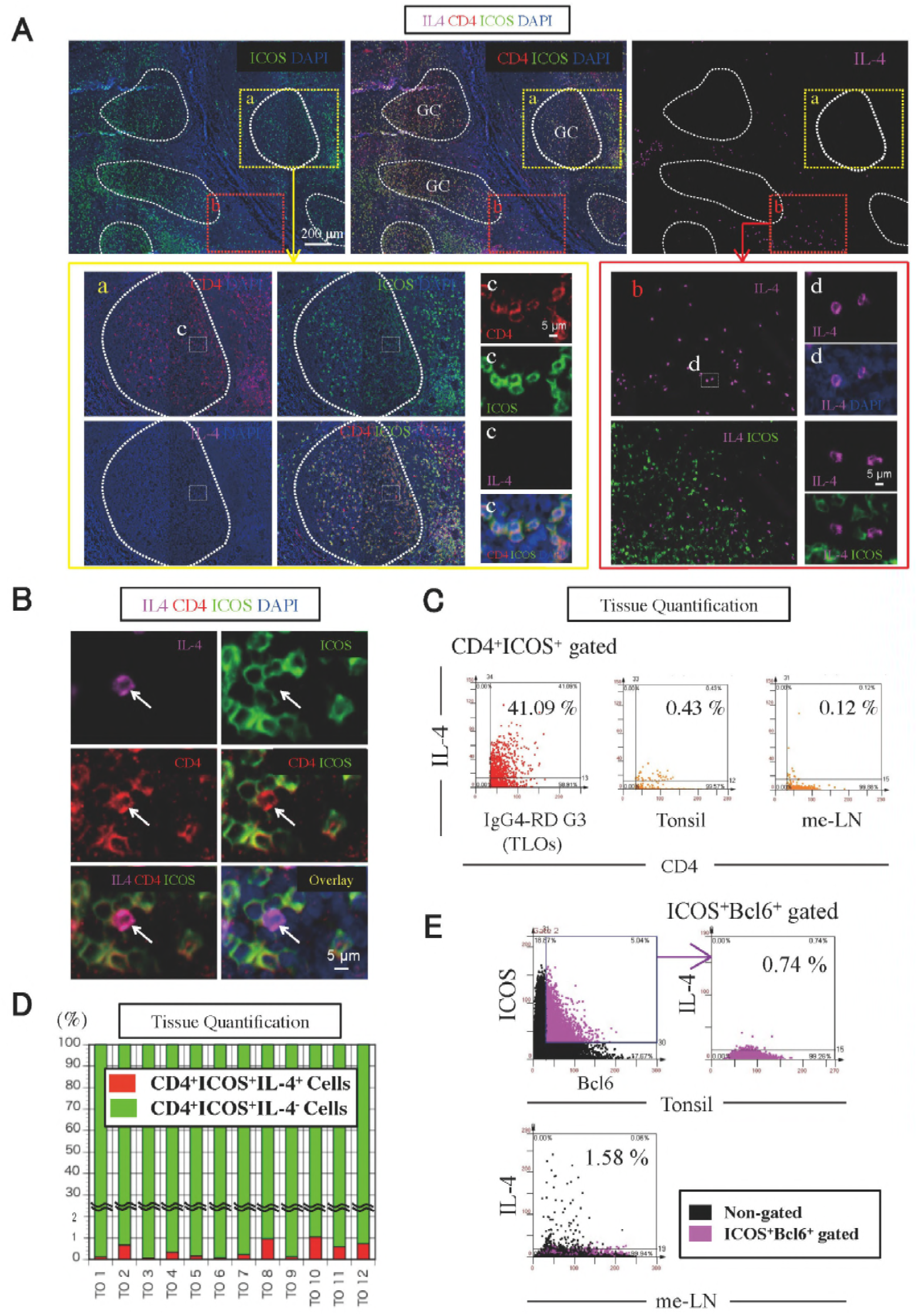
CD4^+^ICOS^+^IL-4^+^ T_FH_ cells are sparse in human secondary lymphoid organs. **(A)** Immunofluorescence staining of CD4 (red), ICOS (green), IL-4 (magenta) and DAPI (blue) in tissues from normal Tonsil. The yellow broken line demarcates the area within germinal centers (GC). The white broken line demarcates the area outside GC. **(B)** Immunofluorescence staining of CD4 (red), ICOS (green), IL-4 (magenta) and DAPI (blue) in tissues from normal tonsils. **(C)** Scatter plots depict the mean fluorescence intensity per cell quantified using TissueQuest software for each fluorescent antibody used to stain tissue from an IgG4-RD subject, normal tonsil, and normal mesenteric lymph node. **(D)** CD4^+^ICOS^+^IL-4^+^ (red) and CD4^+^ICOS^+^IL-4^-^ (green) cells were quantified in tissue from 12 tonsils. **(E)** Scatter plots depict the mean fluorescence intensity per cell quantified using TissueQuest software for each fluorescent antibody used to stain normal tonsil and mesenteric lymph node

In order to more directly examine germinal center T_FH_ cells, we initially localized germinal centers in twelve different human tonsils using staining for Bcl-6 and then analyzed CD4^+^Bcl-6^+^IL-4^+^ T cells (Fig. 2A). These analyses also revealed that CD4^+^Bcl-6^+^IL-4^+^ germinal center T_FH_ cells represented a very small proportion (ranging from 1-10%) of all germinal center T_FH_ cells.

**Fig. 2.**
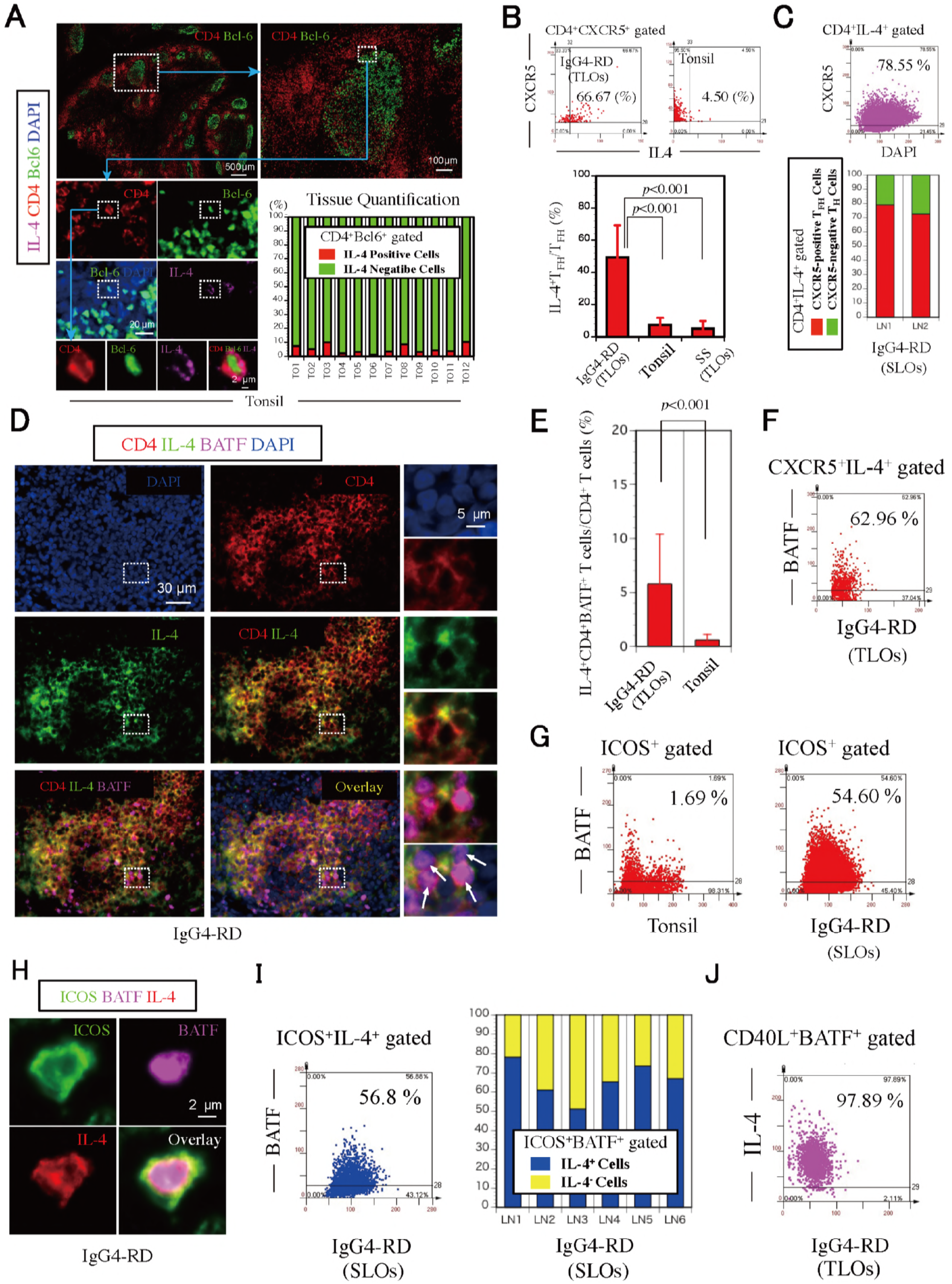
IL-4^+^BATF^+^ TFH cells are rare in normal secondary lymphoid organs but abundant in IgG4-RD. **(A)** Immunofluorescence staining of CD4 (red), Bcl-6 (green), IL-4 (magenta) and DAPI (blue) in normal tonsil. Quantification of CD4^+^Bcl-6^+^IL-4-positive T_FH_ and CD4^+^Bcl-6^+^IL-4-negative T_FH_ in 12 tonsils. **(B)** Scatter plots depict the mean fluorescence intensity per cell quantified using TissueQuest software for each fluorescent immunostain in IgG4-RD SMGs (G1) and normal tonsils. Quantification of CD4^+^CXCR5^+^IL-4^+^ and CD4^+^CXCR5^+^ T_FH_ cells in 17 IgG4-RD SMGs, 12 tonsils, and 7 SS salivary glands. *P* value is based on Mann-Whitney *U* test. **(C)** Almost all IL-4 expressing CD4^+^ T cells in IgG4-RD lymph nodes were CXCR5^+^ TFH cells. Scatter plots depict the mean fluorescence intensity per cell quantified using TissueQuest software for each fluorescent immunostain in IgG4-RD lymph node (LN1). The quantification of CD4^+^IL-4^+^CXCR5-positive T_FH_ and CD4^+^IL-4^+^CXCR5-negative non-T_FH_ cells in lymph nodes from 2 IgG4-RD patients (LN1 and LN2) is shown. **(D)** CD4^+^IL-4^+^BATF^+^ T cells were enriched in IgG4-RD SMGs. Immunofluorescence staining of CD4 (red), IL-4 (green), BATF (magenta) and DAPI (blue) in IgG4-RD SMGs (G12). White arrows indicate CD4^+^IL-4^+^BATF^+^ T cells. **(E)** Quantification of CD4^+^BATF^+^IL-4^+^ T cells and CD4^+^ T cells in 17 IgG4-RD SMGs and 12 normal tonsils. *P* value is based on Mann-Whitney *U* test. **(F)** Scatter plots depict the mean fluorescence intensity per cell quantified using TissueQuest software for each fluorescent antibody used to stain normal tonsil and IgG4-RD lymph node (LN1). **(G)** Scatter plots depict the mean fluorescence intensity per cell quantified using TissueQuest software for each fluorescent immunostain in IgG4-RD SMGs (G3). **(H)** Immunofluorescence staining of ICOS (green), BATF (magenta), and IL-4 (Red) in IgG4-RD LN (LN1). **(I)** Scatter plots depict the mean fluorescence intensity per cell quantified using TissueQuest software for each fluorescent immunostain in IgG4-RD LN (LN1). The quantification of ICOS^+^BATF^+^IL-4-positive T and ICOS^+^BATF^+^IL-4-negative T cells in lymph nodes from 6 IgG4-RD patients (LN1-LN6). **(J)** Scatter plots depict the mean fluorescence intensity per cell quantified using TissueQuest software for each fluorescent immunostain in IgG4-RD SMGs (G3).

### CD4^+^CXCR5^+^IL-4^+^ T_FH_ cells are abundant in IgG4-RD but rare in normal secondary lymphoid organs and in Sj**ö**gren’s syndrome

We analyzed secondary lymphoid organs from healthy individuals and affected tertiary lymphoid organs (TLO) in submandibular glands from IgG4-RD subjects with active disease for IL-4 synthesizing T_FH_ cells and focused our analyses in the vicinity of TLO-like structures containing germinal centers. TLOs with germinal centers (Ruddle, 2014) were identified using multi-color immunofluorescence approaches (CD4, CD19, Bcl6, and DAPI). In this study, 17 of 25 (68%) patients with IgG4-RD and 7 of 15 (47%) patients with severe Sjögren’s syndrome (SS) had TLOs in affected salivary glands (Table S3). Germinal centers within TLOs were both more frequent and larger in salivary glands from IgG4-RD than severe SS (Table S3). As seen in Fig. 2B, CD4^+^CXCR5^+^IL-4^+^ cells were found in human tonsils and lymph nodes, but were sparse. In contrast, these CD4^+^CXCR5^+^IL-4^+^ T_FH_ cells were abundant in IgG4-RD. Quantitation of CD4^+^CXCR5^+^IL-4^+^ T_FH_ cells revealed that about 67% of CD4^+^CXCR5^+^ T_FH_ cells in an IgG4-RD subject expressed IL-4, while fewer than 5% of tonsillar CD4^+^CXCR5^+^ T_FH_ cells expressed IL-4 (Fig. 2B). CD4^+^CXCR5^+^IL-4^+^ T_FH_ and CD4^+^ICOS^+^IL-4^+^ T_FH_ cells are clearly far more abundant in IgG4-RD than in normal secondary lymphoid organs in or around TLOs from affected tissues from subjects with SS (Fig. 2B).

We extended our studies to lymph nodes from subjects with IgG4-RD. Quantification revealed that almost all IL-4 expressing CD4^+^ T cells in IgG4-RD lymph nodes were CXCR5^+^ T_FH_ cells (Fig. 2C).

### IL-4^+^BATF^+^ T_FH_ cells are rare in normal secondary lymphoid organs but abundant in IgG4-RD

The regulation of IL-4 expression in murine T_FH_ cells is distinct from the mechanisms that drive IL-4 expression in T_H2_ cells. BATF is a transcription factor that may regulate IL-4 secretion by murine T_FH_ cells (Sahoo et al., 2015), but may more broadly help identify T_FH_ cells, some of which do not necessarily express IL-4 (Weinstein et al., 2016). Many CD4^+^IL-4^+^ T cells in the vicinity of germinal centers in affected IgG4-RD tissues express nuclear BATF (Fig. 2D). Not surprisingly, quantification revealed that these IL-4 expressing CD4^+^BATF^+^ T cells are abundant in IgG4-RD tissue lesions but rare in normal SLOs (Fig. 2E). Using parallel sections of tissues from an IgG4-RD subject, we noted that CD4^+^BATF^+^IL-4^+^ T cells were located in the same region in which IL-4 expressing CD4^+^CXCR5^+^T_FH_ cells were observed. Quantification of CXCR5^+^BATF^+^IL-4^+^ cells revealed that about 60% of CXCR5^+^IL-4^+^ cells in an IgG4-RD subject with TLOs expressed BATF (Fig. 2F).

We considered the possibility that the expansion of these IL-4 expressing T_FH_ cells might represent an important disease-related T_FH_ subset and contribute to the specific class switch event in IgG4-RD. ICOS^+^BATF^+^ T_FH_ cells are far more abundant in IgG4-RD lymph nodes than in normal tonsils (Fig. 2G). As seen in Figure 2H, ICOS^+^BATF^+^IL-4^+^ T_FH_ cells were detected in IgG4-RD patients, and these cells were abundant. Quantification of ICOS^+^BATF^+^IL-4^+^ T_FH_ cells revealed that about 60% of ICOS^+^IL-4^+^ T_FH_ cells in an IgG4-RD lymph node subject expressed BATF (Fig. 2I). Furthermore, quantification revealed that the majority of ICOS^+^BATF^+^ T_FH_ cells in IgG4-RD lymph nodes expressed IL-4 as well (Fig. 2I). Interestingly, quantification of CD40L^+^BATF^+^IL-4^+^ T cells revealed that these cells represented approximately 98% of CD40L^+^BATF^+^ T cells in an IgG4-RD subject (Fig. 2J). These data indicate that a large number of lesional IL-4^+^ T_FH_ cells in IgG4-RD express BATF, CD40L and ICOS.

### IL-4 secreting T_FH_ cells are a distinct population of T follicular helper cells

Although visualization by multicolor staining permits anatomic localization of T_FH_ cells in tissues, it can provide only limited information about a few expressed proteins in any putative T_FH_ cell subset, and a detailed characterization of any specific cytokine secreting T_FH_ subset is currently lacking. In order to better understand the biology of IL-4 secreting T_FH_ cells found in lymphoid organs, we performed RNA-seq analysis of viable IL-4 secreting T_FH_ cells from human tonsils. Although the fraction of IL-4 producing T_FH_ cells is low in human tonsils, we were able to purify this subset by starting with 600 million tonsil cells, using an IL-4 cytokine capture strategy. To promote cytokine secretion, CD19-depleted lymphocytes from tonsils were stimulated overnight with plate-coated anti-CD3 and anti-CD28 antibodies. We then compared the transcriptome of FACS sorted IL-4 producing T_FH_ cells to CXCR5^hi^PD1^hi^ T_FH_ cells that did not secrete IL-4, as well as IL-4-secreting CD45RA^-^CXCR5^-^ non-T_FH_ cells obtained from the same tonsil (Figure 3A). CD4^+^ CD45RA^+^ naïve cells were also included as an additional control. Of the 26,002 mapped transcripts, 7,792 were differentially expressed across the four conditions (Figure 3B). The IL-4 producing and non-producing T_FH_ cells were most similar, and markedly dissimilar from IL4-secreting non-T_FH_ cells or naïve CD4^+^ T cells (Figure 3C). In contrast to the IL-4 producing non-T_FH_ cells that express high levels of IL-4, IL-5 and IL-13, reflecting a T_H2_ signature, IL-4 producing T_FH_ cells express only IL-4 but much lower levels of IL-5 and IL-13 (Figure 3D). In order to examine lineage-defining genetic regulators and functionally relevant effector molecules, we focused our analysis on differentially expressed CD molecules, transcription factors and cytokines (Figure 3E). The transcript level of genes critical to T_FH_ function such as CD40L, ICOS, CXCR5, IL-21R and PD1 was highest in the IL4-secreting T_FH_ cells. In addition, they also expressed high levels of CCR4, CD200, CTLA4, GITR and low levels of CD6, CD27, CD28, SELL, IL-7R and CD74. Although the expression levels of these cell surface markers are derived from *in vitro* activation during cytokine capture, the CD markers specific for cells with an IL-4 secreting T_FH_ phenotype may aid in the specific flow cytometric identification of *ex vivo* human IL-4 secreting T_FH_ cells in future studies. As expected, CD4^+^ CD45RA^+^ cells had the highest level of CCR7, CD27 and CD62L. The high expression levels of multiple chemokines, cytokines and their receptors including CCR2, CCR6, CXCR3, CXCR6, IL-2RA, IL-2RB, IL-10RA, CCL4, IFNG, IL-2, IL-3, IL-4, IL-5, IL-9, IL-10, IL-17A, IL-22 and IL-23A within the IL4-producing CD4^+^CD45RA^-^ non-T_FH_ cells perhaps indicate the heterogeneity among these cells, and may reflect the overlap and plasticity between T_H1_, T_H2_ and T_H17_ subsets that is seen following TCR stimulation in the absence of polarizing cytokines (i.e., anti-CD3 and anti-CD28 alone). Interestingly, the IL-4 producing T_FH_ cells express the highest levels of BCL6 and BATF in this comparison, but not transcription factors related to T_H2_ differentiation such as GATA3, STAT5A and PRDM1 (BLIMP1), which are instead abundantly expressed in the IL-4-producing non-T_FH_ population, which is likely enriched for T_H2_ cells. Furthermore, the expression of all the markers that we studied using immunofluorescence in IL-4 expressing T_FH_ cells from IgG4-RD tissues (BCL6, ICOS, IL-4, CXCR5, BATF, GATA3 and PD1) were consistent with the RNA-seq observations on the tonsillar IL-4 secreting T_FH_ cells.

**Fig 3.**
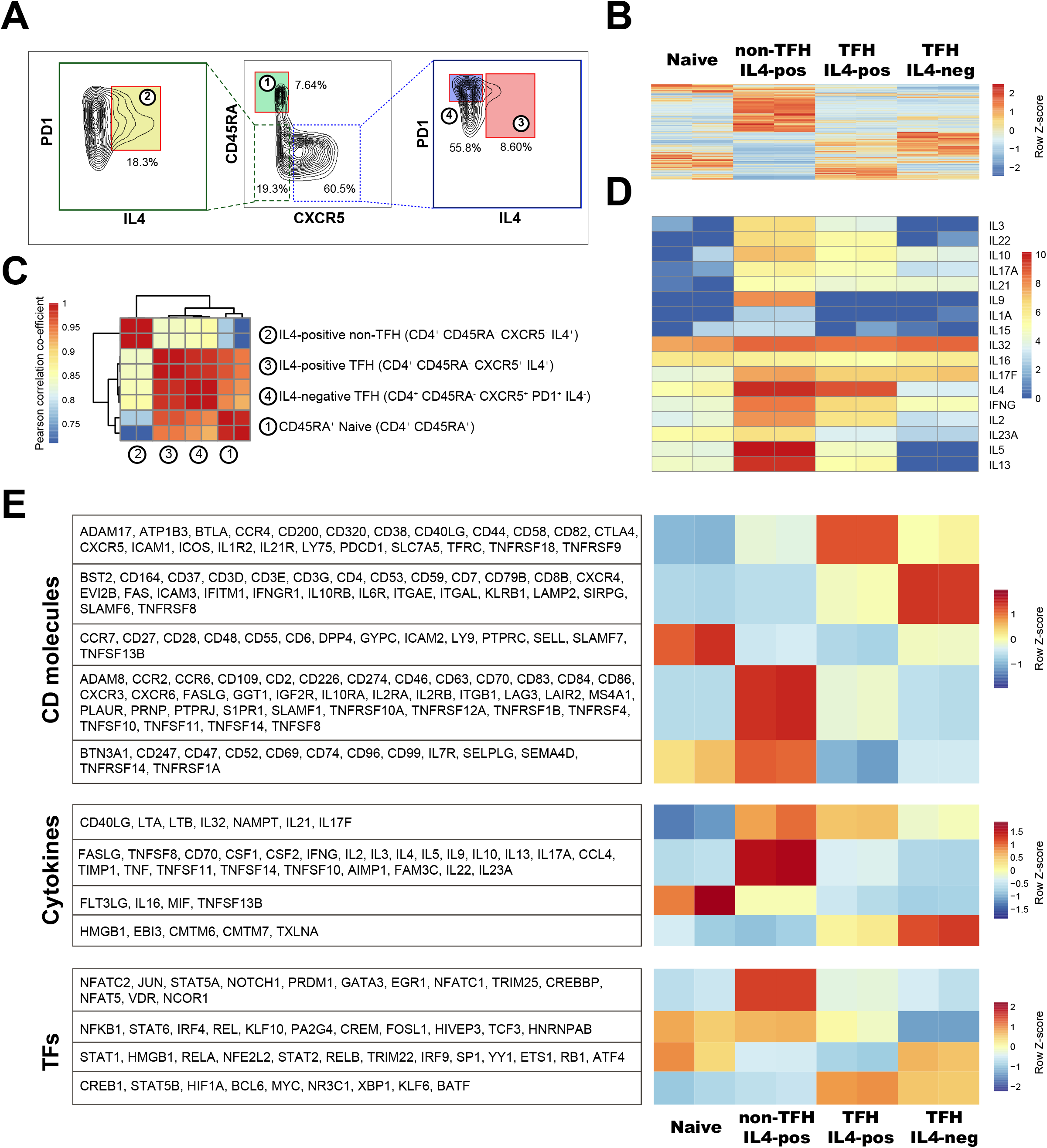
Transcriptomic profiling of IL-4 producing T_FH_ cells from human tonsil. **(A)** Gating strategy used to sort IL-4 secreting and non-secreting tonsillar CD45RA^+^CXCR5^+^ T_FH_ cells following anti-CD3/anti-CD28 stimulation. CD45RA^+^ CXCR5^-^ cells, and CD45RA^-^CXCR5^-^IL4^+^ cells were sorted as additional controls. **(B)** A heatmap of differentially expressed genes across all four conditions depicting the Z-scores for normalized expected read counts. **(C)** A correlation matrix of differentially expressed genes. **(D)** Expression pattern of individual interleukins is depicted on a log scale. **(E)** Expression of CD molecules, cytokines and transcription factors clustered into patterns using k-means. Z-scores of the expected read counts for each cluster are shown.

### CD4^+^CXCR5^+^IL-4^+^ T_FH_ cells are mainly outside germinal centers in IgG4-RD and sometimes physically associate with AID expressing B cells

As seen in Figure 4A, CD4^+^CXCR5^+^IL-4^+^ T_FH_ cells were abundant in affected IgG4-RD tissues and were mainly outside germinal centers, but were also located in the light zone within GCs. Using parallel tissue-sections, we noted that IgG4-positive B cells were abundant outside germinal centers in the same region in which IL-4 secreting T_FH_ cells were observed. We quantitated IL-4 expressing T_FH_ cells and IgG4-expressing B cells within and outside germinal centers and observed that the majority of both cell types reside outside germinal centers (Figs. 4A and 4B).

**Fig. 4.**
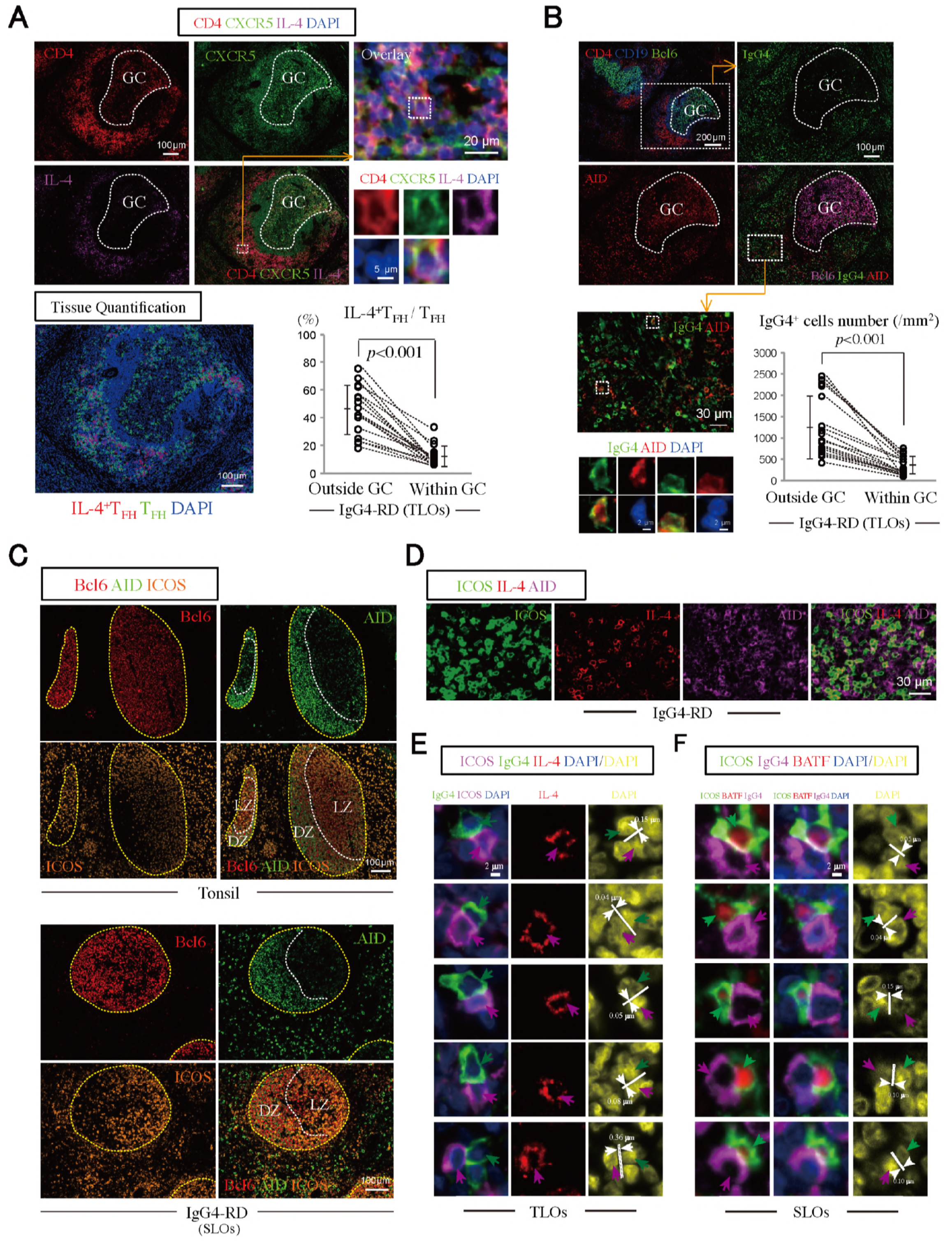
CD4^+^CXCR5^+^IL-4^+^ T_FH_ cells are abundant around GCs in IgG4-RD and sometimes physically contact AID expressing B cells. **(A)** CD4^+^CXCR5^+^IL-4^+^ T cells were enriched around TLOs in IgG4-RD SMGs. Immunofluorescence staining of CD4 (red), IL-4 (magenta), CXCR5 (green) and DAPI (blue) in IgG4-RD SMGs (patient G3). Quantification of CD4^+^CXCR5^+^IL-4^+^ T_FH_ cells and CD4^+^CXCR5^+^ T_FH_ cells comparing those outside and within germinal centers from each of five different areas in TLOs of 17 IgG4-RD patients (G1-G17). *P* value is based on Mann-Whitney *U* test. **(B)** IgG4^+^B cells were enriched outside GC, especially some these cells express AID. Immunofluorescence staining of CD4 (red), CD19 (blue), and Bcl6 (green) in IgG4-RD SMGs (patient G3). Immunofluorescence staining of AID (red), Bcl6 (magenta), IgG4 (green) and DAPI (blue) in IgG4-RD SMGs (patient G3). The numbers of IgG4^+^ cells (/mm^2^) outside and within germinal centers were quantified from each of five different areas in TLOs of 17 IgG4-RD patients (G1-G17). *P* value is based on Mann-Whitney *U* test. **(C)** AID expressing B cells outside GCs in IgG4-RD lymph node were abundant. Immunofluorescence staining of Bcl6 (red), AID (green), and ICOS (orange) in a control tonsil and IgG4-RD lymph node. LZ, light zone; DZ, dark zone. **(D)** Immunofluorescence staining of ICOS (green), IL-4 (Red), and AID (magenta) in IgG4-RD SMGs. **(E)** Immunofluorescence staining of IgG4 (green), ICOS (magenta), IL-4 (red) and DAPI (blue) in TLOs with an IgG4-RD subject (G3). Magenta arrows indicate ICOS^+^IL-4^+^ T_FH_ cells. Green arrows indicate IgG4^+^ B cells. A number of IL-4 expressing T_FH_ cells and IgG4^+^ B cells formed close and extensive inter-cellular plasma membrane contacts. The distance was measured between the edge of each IgG4^+^ B cell nucleus to the edge of the closest IL-4 expressing TFH cell nucleus, and an inter-nuclear distance of less than 0.4 µm was indicative of a T-B conjugate. **(F)** Immunofluorescence staining of ICOS (green), IgG4 (magenta), BATF (red) and DAPI (blue) in lymph nodes of an IgG4-RD subject. Green arrows indicate ICOS^+^BATF^+^ T_FH_ cells. Magenta arrows indicate IgG4^+^ B cells. A number of BATF^+^ICOS^+^ T_FH_ cells and IgG4^+^ B cells formed close and extensive inter-cellular plasma membrane contacts. The distance was measured between the edge of each IgG4^+^ B cell nucleus to the edge of the closest BATF^+^ICOS^+^ TFH cell nucleus. All inter-nuclear distances in this figure were less than 0.2 µm.

It is generally accepted that CD40L-CD40 signaling induces activation induced cytidine deaminase (AID) and that specific cytokines target selected switch regions. Do the IL-4 expressing T_FH_ cells and AID expressing B cells make physical contact with one another? We examined lymph nodes and TLOs from IgG4-RD subjects to determine whether ICOS^+^ T_FH_ cells are in physical contact with AID expressing B cells or IgG4 expressing B cells *in situ*. AID expressing B cells could be visualized outside germinal centers in IgG4-RD sections (Figs. 4B and 4C); we noted that IgG4 expressing B cells outside germinal centers often express AID (Fig. 4B). We also noted the existence of IL-4^+^ICOS^+^ T_FH_ cells in cell-cell contact with AID expressing B cells (Fig. 4D). As has been reported earlier, much of the AID staining seen is cytosolic (Cattoretti et al., 2006). Furthermore, we noted the existence of IL-4^+^ICOS^+^ and BATF^+^ICOS^+^ T_FH_ cells that are in cell-cell contact with IgG4 expressing B cells (Fig. 4E and 4F), confirming visual contact with nuclear distance measurements. AID^+^IgG4^+^ B cells were also visualized within and outside GCs in TLOs in IgG4-RD.

### CD4^+^CXCR5^+^IL-4^+^ T_FH_ cells and CD4^+^IL-4^+^BATF^+^ T cells are enriched in IgG4-RD, and their proportions are tightly linked to serum IgG4 levels and the proportion of IgG4 positive plasma cells in tissues

As seen in Fig. 5A, only a small proportion of CD4^+^CXCR5^+^ T_FH_ cells in tonsils, mesenteric and cervical lymph nodes from normal individuals and also in or around TLOs from affected tissues from subjects with SS synthesize IL-4. These data argue that IL-4 expression is not generally a part of the T_FH_ phenotype in cells located in secondary or tertiary lymphoid organs. Instead, what is more likely is that a subset of activated T_FH_ cells that expresses IL-4 expands considerably in a disease in which there is a prominent IgG4 class switch, but these cells are not abundant around GCs in healthy people or in TLOs from other diseases in which there is no prominent switch to the IgG4 or IgE isotypes. The proportion of T_FH_ cells that express IL-4 in disease tissue correlates very strongly with the serum IgG4 levels in IgG4-RD subjects, but not with total serum IgG, IgE or IgA levels (Fig. 5B). A negative correlation with serum IgM levels was also observed (Fig. 5B). The proportion of CD4^+^CXCR5^+^ cells that express IL-4 in IgG4-RD also correlates with the proportion of IgG4 expressing plasma cells in disease tissues (Fig. 5C). The proportions of CD4^+^BATF^+^IL-4^+^ T cells correlate with CD4^+^CXCR5^+^IL-4^+^ T_FH_ cells and with serum IgG4 levels (Fig. 5D).

**Fig. 5.**
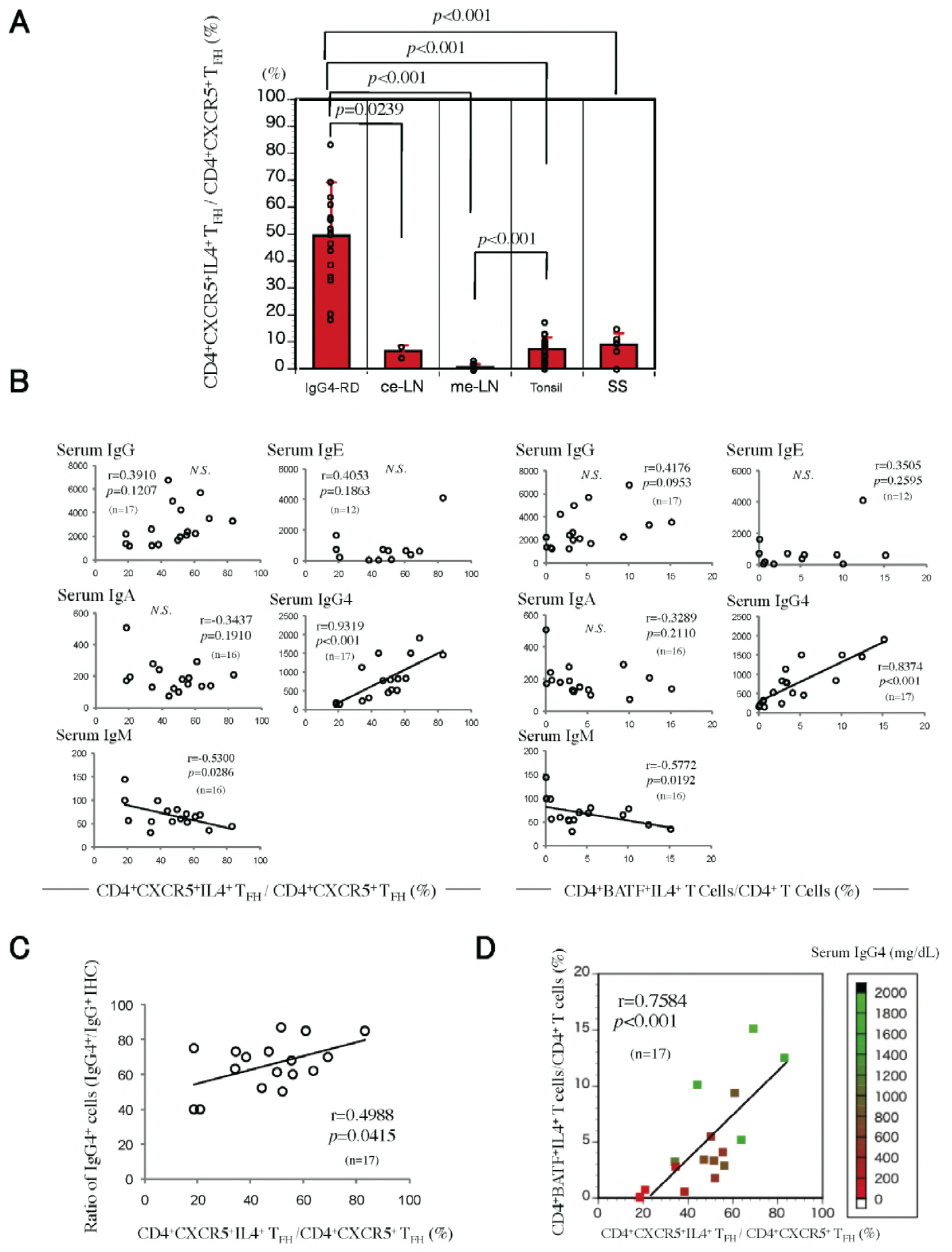
CD4^+^CXCR5^+^IL-4^+^ T_FH_ cells and CD4^+^IL-4^+^BATF^+^ T cells are enriched in IgG4-RD and their proportions are tightly linked to serum IgG4 levels and the proportion of IgG4 positive plasma cells. **(A)** Quantification of CD4^+^CXCR5^+^IL-4^+^ and CD4^+^CXCR5^+^ T_FH_ cells in 17 IgG4-RD SMGs, 7 SS LSGs, 12 tonsils, 2 cervical lymph nodes and 3 mesenteric lymph nodes. *P* value is based on Mann-Whitney *U* test. **(B)** Correlations of the proportion of CD4^+^CXCR5^+^IL-4^+^ T_FH_ cells and CD4^+^BATF^+^IL-4^+^ T cells in SMGs from IgG4-RD patients and their serum IgG, IgA, IgE, IgM or IgG4 levels. The correlation coefficients and *P* value were determined using Spearman’s rank correlations. **(C)** Correlations of the proportions of CD4^+^CXCR5^+^IL-4^+^ T_FH_ cells in SMGs from IgG4-RD patients and their ratio of IgG4^+^/IgG^+^ cells (n=17). The correlation coefficients and *P* value were determined using Spearman’s rank correlations. **(D)** The frequency of CD4^+^CXCR5^+^IL-4^+^ T_FH_ cells in SMGs from IgG4-RD patients (n=17) correlated with the frequency of CD4^+^BATF^+^IL-4^+^ T cells and serum IgG4 concentrations. The correlation coefficients and *P* value were determined using Spearman’s rank correlations.

## DISCUSSION

In T-dependent immune responses, it is widely accepted that specific cytokines induce the transcription of selected switch regions so that distinct Ig gene loci are targeted by AID for the induction of cytidine deamination and double strand break formation. However, a link between subsets of T_FH_ cells and specific isotype switching events has not been established.

We found that very few T_FH_ cells in healthy tonsils or lymph nodes synthesize IL-4. These findings are broadly consistent with published data on human tonsillar T_FH_ cells using different approaches (Bentebibel et al., 2011; Kroenke et al., 2012; Ma et al., 2009). Bentebibel et al. reported that T_FH_ cells from outside and within GCs secrete IL-4 when co-cultured *in vitro* with B cells. However, as these data did not provide information at the single cell level, the percentage of T_FH_ cells capable of secreting IL-4 could not be surmised from this report. Kroenke et al. used flow cytometry to quantitate intracellular cytokines in *in vitro* re-stimulated T_FH_ cells from tonsils. Following re-stimulation with PMA and ionomycin, less than 3% of tonsillar T_FH_ cells expressed IL-4 (Kroenke et al., 2012). Ma et al. also used flow cytometry to measure cytokine expression by tonsillar T_FH_ cells following PMA and ionomycin exposure and observed that up to 16% of CXCR5^hi^ T_FH_ cells expressed IL-4 (Ma et al., 2009). Our *in situ* data, based on a direct examination of IL-4 in T_FH_ cells, rather than on *in vitro* re-stimulation, indicates that only a small fraction of T_FH_ cells in normal human SLOs produce IL-4. These data suggest that IL-4 secreting T_FH_ cells are relatively rare in human tonsils to begin with, and only a small fraction of healthy tonsillar T_FH_ cells have presumably been re-stimulated *in vivo* to synthesize IL-4, as evidenced by our *in situ* studies. In IgG4-RD, in contrast, a very large proportion of T_FH_ cells that can secrete IL-4 infiltrate tissues and secondary lymphoid organs, and because these cells have likely been re-stimulated *in vivo*, they can be seen to express IL-4 in our *in situ* analyses.

Human T_FH_ subsets have only been described in the context of putative circulating memory CD4^+^ T cells and not in a tissue context. Studies on T_FH_ cells in disease have generally focused on an analysis of the blood. We examined human T_FH_ cells *in situ* using multi-color quantitative immunofluorescence approaches, focusing on lymph nodes and tertiary lymphoid organs in tissues of subjects with a disease in which there is a pronounced switch to the IgG4 isotype. The proportions of CD4^+^CXCR5^+^IL-4^+^ T cells, and their likely equivalent, CD4^+^BATF^+^IL-4^+^ T cells, are markedly increased in IgG4-RD (with most IL-4 expressing T_FH_ cells located in lymphoid cuffs just outside germinal centers), and these proportions correlate well with serum IgG4 levels and the proportion of IgG4-positive cells in disease tissues. Our data links IL-4 expressing T_FH_ cells to the IgG4 class switch and also indirectly argues that most class switching likely occurs outside germinal centers.

IL-4 secreting T_FH_ cells have never been purified from lymphoid organs and cannot be distinguished in human lymphoid organs by the expression of specific chemokine receptors. To reliably characterize a specific cytokine-secreting T_FH_ population, we used a specific IL-4 capture assay and compared the transcriptomes of IL-4 producing human tonsillar T_FH_ cells to non-T_FH_ cells from the tonsil that secrete IL-4, and human tonsillar PD-1^hi^ T_FH_ cells that do not secrete IL-4. IL-4 secreting T_FH_ cells stand out as a distinct subset. It is reasonable to consider that these IL-4 secreting BATF expressing T_FH_ cells, which do not express GATA-3, represent an activated T_FH_ population or subset in secondary and tertiary lymphoid organs that are expanded in disease that may contribute to class switching to IgG4. Unlike T_H2_ cells, which produce IL-4, IL-5 and IL-13, the IL4-producing tonsillar T_FH_ cells produce IL-4 and IL-10, but not IL-5 or IL-13. This subset of T_FH_ cells has a unique transcriptional profile and a distinct set of surface markers that will facilitate future functional studies.

The mechanisms and signals by which antibodies class switch to IgG4 is poorly understood. CD4^+^ T cells can activate IgM-positive B cells to switch to IgG4 and IgE in the presence of added IL-4 (Gascan et al., 1991). It has been argued based on *in vitro* studies that IL-10 may contribute to the IgG4 class switch indirectly by somehow facilitating IL-4 mediated switching to IgG4 in preference to IgE (Jeannin et al., 1998). No molecular or cellular explanation has emerged for this *in vitro* phenomenon. Arguments have also been made that switching to IgG4 is a result of repeated division, and that switching progresses sequentially along chromosome 14 from IgG1 to IgG3 to IgG2 to IgG4 (Tangye et al., 2002) (Jackson et al., 2014). We do not presume that IL-4 secreting T_FH_ cells alone contribute to the IgG4 class switch, though it is possible that the same subset of IL-4 secreting T_FH_ cells that we have characterized might, in certain contexts, make enough IL-10 to facilitate this switching event. Perhaps more efficient purification of this T_FH_ subset using surface markers inferred from our transcriptomic data will facilitate future functional studies on IL-4 secreting T_FH_ cells. Tissue sources of IL-10 or of other factors that work in concert with IL-4 from these specific T_FH_ cells will also need to be explored in future studies.

These studies are the first demonstration of an *in situ* expansion of what may be construed to be a human T_FH_ subset. Many diseases exist in which IgE dominates in the absence of IgG4 and *vice versa*. Our data indicate that only a fraction of T_FH_ cells in secondary and tertiary lymphoid organs, mainly outside germinal centers, express BATF and IL-4. However, we have shown here that these IL-4 expressing T_FH_ cells accumulate in tertiary lymphoid organs of subjects with IgG4-RD. There is therefore a disease-specific increase in the accumulation of a subset of T_FH_ cells and a tight association between the numbers of these cells and class switching, specifically to IgG4. Whether there are further subsets of BATF and IL-4 expressing T_FH_ cells that separately facilitate the IgG4 or IgE class switch remains to be ascertained using single cell approaches.

## METHODS

### Study population

Submandibular glands (SMGs) were obtained from 17 Japanese patients with IgG4-RD, from affected lymph nodes of 6 patients with IgG4-RD, and from lacrimal and salivary glands (LSGs) of 7 patients with active Sjögren’s Syndrome (SS). In addition, 2 unaffected cervical lymph nodes, 3 mesenteric lymph nodes and 12 healthy human tonsils were obtained that were histologically normal. All of these IgG4-RD patients had been followed up between 2007 and 2015 at the Department of Oral and Maxillofacial Surgery of Kyushu University Hospital, a tertiary care center. Open SMG biopsies were obtained from patients with IgG4-RD (Moriyama et al., 2014). IgG4-RD was diagnosed according to the following criteria (Umehara et al., 2012): 1) persistent (longer than 3 months) symmetrical swelling of more than two lacrimal and major salivary glands; 2) high (>135 mg/dl) serum concentrations of IgG4; and 3) infiltration of IgG4-positive plasma cells into tissue (IgG4^+^ cells/IgG^+^ cells >40%), as determined by immunostaining. All SMGs from patients with IgG4-RD had histopathologic features of IgG4-RD. The age, sex, serum immunoglobulin and specific autoantibody levels of 17 IgG4-RD patients (G1-G17) whose affected salivary gland biopsies were analyzed by *ex vivo in situ* immunofluorescence studies are summarized in Table S1A. The age, sex, serum immunoglobulin and specific autoantibody levels of 6 IgG4-RD patients (LN1-LN6) whose affected lymph node biopsies were analyzed by *ex vivo in situ* immunofluorescence studies are summarized in Table S2.

Each SS patient exhibited objective evidence of salivary gland involvement based on the presence of subjective xerostomia and a decreased salivary flow rate, abnormal findings on parotid sialography and focal lymphocytic infiltrates in the LSGs by histology (Vitali et al., 2002). All SS patients were severe cases and had developed abundant TLOs in each salivary gland. None of the IgG4-RD and SS patients had a history of treatment with steroids or other immunosuppressants, infection with HIV, hepatitis B virus, or hepatitis C virus, or sarcoidosis, and none had evidence of malignant lymphoma at the time of the study. The age, sex, serum immunoglobulin and specific autoantibody levels of SS patients whose salivary gland biopsies were analyzed by *in situ* immunofluorescence are summarized in Table S1B.

Normal human mesenteric lymph node and tonsil subjects were obtained from Massachusetts General Hospital. Normal human cervical lymph nodes were obtained from the Department of Oral and Maxillofacial Surgery of Kyushu University Hospital.

The study protocol was approved by the Ethics Committee of Kyushu University, Japan and the Institutional Review Board at Massachusetts General Hospital. All included disease subjects provided written informed consent.

### Multi-color immunofluorescence staining

Tissue samples were fixed in formalin, embedded in paraffin, and sectioned. These specimens were incubated with antibodies: anti-AID (clone: ZA001; Invitrogen), anti-IgG4 (clone: ab109493; Abcam) anti-ICOS (clone: 89601 Cell Signaling Technologies), anti-IL-4 (clone: MAB304; R&D Systems), GATA3 (clone: CM405A; Biocare), CXCR5 (clone: MAB190; R&D), Bcl-6 (clone: CM410A,C; Biocare), BATF (clone: 10538; Cell Signaling), CD4 (clone: CM153A; Biocare) and CD19 (clone: CM310 A,B; Biocare) followed by incubation with secondary antibody using a SuperPicTure™ Polymer Detection Kit (Invitrogen) and an Opal™ 3-Plex Kit (Fluorescein, Cyanine3, and Cyanine5). The samples were mounted with ProLong^TM^ Gold Antifade mountant containing DAPI (Invitrogen).

### Microscopy and quantitative image analysis

Images of the salivary gland specimens were acquired using the TissueFAXS platform (TissueGnostics). For quantitative analysis, the entire area of the tissue involved by the lymphoplasmacytic infiltrate was acquired as digital greyscale images in four channels with filter settings for FITC, Cy3 and Cy5 in addition to DAPI. Cells of a given phenotype were identified and quantitated using the TissueQuest software (TissueGnostics), with cut off values determined relative to the positive controls. This microscopy-based multicolor tissue cytometry software permits multicolor analysis of single cells within tissue sections similar to flow cytometry. The principle of the method and the algorithms used have been described in detail elsewhere (Ecker and Steiner, 2004).

### Evaluation of TLOs with IgG4-RD and SS

TLOs with germinal centers (Ruddle, 2014) were identified by using multi-color immunofluorescence approaches (CD4, CD19, Bcl6, and DAPI). In this study, SMG and LSG tissue sections from 25 patients with IgG4-RD and 15 patients with severe SS were evaluated. Distinct Bcl-6^+^ germinal centers were observed in affected IgG4-RD tissues. Bcl-6^+^ germinal center B cells in TLOs were within B cell follicles, delineated using antibodies to CD19. Seventeen of 25 (68%) patients with IgG4-RD had TLOs in affected salivary glands. The age, sex, serum immunoglobulin and specific autoantibody levels of 17 IgG4-RD patients (G1-G17) whose affected salivary gland biopsies were analyzed by *in situ* immunofluorescence are summarized in Table S1A. Seven of 15 (47%) patients with severe SS had TLOs in affected salivary glands. The age, sex, serum immunoglobulin and specific autoantibody levels of SS patients whose salivary gland biopsies were analyzed in this study are summarized in Table SS1B. The number of TLOs with germinal centers and the size of germinal centers in each TLOs was evaluated in 4mm^2^ sections from five different areas of 17 IgG4-RD patients (G1-G17) and 7 SS patients (SS1-SS7) (Table S3).

### IL-4 capture assay and cell sorting

600 million cells from human tonsil were re-suspended in Miltenyi MACS buffer and stained with biotinylated anti-human CD19 (clone HIB19, Biolegend) on ice for 25 minutes. Cells were washed once in MACS buffer and incubated with anti-biotin microbeads (Miltenyi Biotec) for 25 minutes on ice. Cells were then loaded on two separate Miltenyi LS columns (at 300 million cells per column) and the flow-through was collected as the B cell depleted fraction. These cells were spun down and stimulated overnight with plate coated anti-CD3 (5µg/ml) and anti-CD28 (5µg/ml). The stimulated cells were then enriched for IL-4-secreting cells using the vendor’s protocol (Miltenyi Biotec IL-4 Secretion Assay kit, #130-054-101). The enriched IL-4+ cell fraction was surface stained with antibodies against CD4 (BioLegend #317420), CD45RA (clone H100, BioLegend #304122), CXCR5 (clone J252D4, BioLegend #356920), and PD1 (clone EH12.2H7, BioLegend #329924). The following populations were viably sorted using FACSAria2 (Becton Dickinson) directly into Qiagen RLT-plus buffer (with 1% 2-mercaptoethanol): (i) IL4-secreting CD4+CXCR5+ TFH cells, (ii) IL4-secreting CD4+CD45RA-CXCR5-Th2 cells, (iii) IL4-negative CD4+CXCR5+PD1+ TFH cells, and (iv) IL4-negative CD45RA+ cells.

### Trancriptomic analyses

Total RNA was isolated from the FACS-sorted cells using the RNeasy plus Micro Kit (Qiagen, USA). RNA sequencing (RNA-Seq) libraries were prepared as previously described (Picelli. et al., 2013). Briefly, whole transcriptome amplification (WTA) and tagmentation-based library preparation were performed using the SMART-Seq2 protocol, followed by 35-base pair paired-end sequencing on a NextSeq 500 instrument (Illumina). 5-10 million reads were obtained from each sample and aligned to the UCSC hg38 transcriptome. Gene expression was calculated using RSEM as previously described (Li and Dewey, 2011). The EBSeq package was used to identify differentially expressed genes with a posterior probability of differential expression (PPDE) > 0.95 (Leng et al., 2013). Our analysis focused on cytokines, transcription factors and CD molecules. Cytokines and transcription factors were obtained using the gene ontology terms GO:0003700 and GO:0005125, respectively. Transcription factors pertaining to immune cells were filtered using a literature-based gene prioritization approach with the query terms “immune response”, “immunity”, “T cells” and “lymphocytes” (Fontaine et al., 2011). The data discussed in this publication have been deposited in NCBI’s Gene Expression Omnibus (Edgar et al., 2002) and are accessible through GEO Series accession number GSE111968 (https://www.ncbi.nlm.nih.gov/geo/query/acc.cgi?acc=GSE111968)

### Statistical analyses

Differences between groups were determined by the *X*^2^ test, Student t test, Mann–Whitney *U* test and Spearman rank correlations. All statistical analyses were performed using JMP Pro software, version 11 (SAS Institute, Cary, NC, USA) for Mac. *P* values less than 0.05 were considered statistically significant. Nonsignificant differences were not specified. In all figures, bar charts and error bars represent the means ± SEM, respectively.

## ACKNOWLEDGEMENTS

This work was supported by awards AI110495 (to SP) and AI113163 (to VSM) from the National Institute of Health and supported by the Japanese Society for the Promotion of Science (JSPS) Postdoctoral Fellowships for Research Abroad to TM and Mochida Memorial Medical and Pharmaceutical Foundation to TM. The authors thank Eric Safai for help with the collection and processing of samples from Massachusetts General Hospital and Thomas Diefenbach of the Imaging Core at the Ragon Institute for help and advice.

## Supplemental items

**Table S1.**
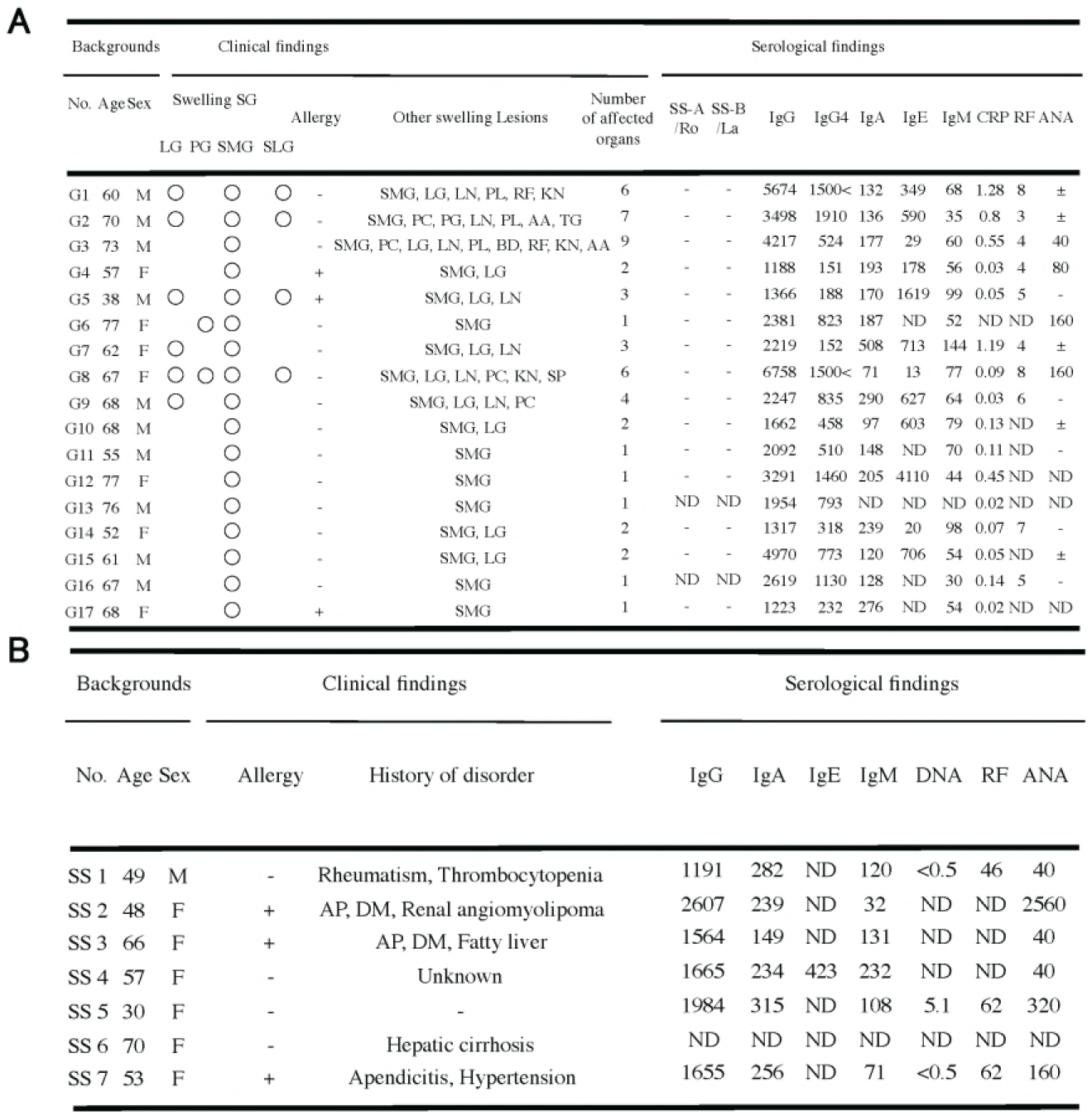
Background information, Clinical and Serological findings for 17 IgG4-RD and 7 SS patients whose affected salivary gland biopsies were analyzed in *ex vivo in situ* immunofluorescence studies. Abbreviations: SG, salivary gland; LG, lacrimal gland; PG, parotid gland; SMG, salivary mandibular gland; SLG, sublingual gland; LN, lymph node; BD, bile duct; PC, pancreas; PS, prostate; AA, aorta abdominalis; LG, lacrimal gland; RF, retroperitoneal fibrosis; KN, kidney; PL, pleura; PG, pituitary gland; TG, thyroid gland; SP, spleen; OM, orbital muscle; AP, Autoimmune pancreatitis; DM, Diabetes; -, negative; ND, not done.

**Table S2.** Background information and Serological findings for 6 IgG4-RD patients whose affected lymph node biopsies were analyzed by *ex vivo in situ* immunofluorescence studies. ND, not done.

| Backgrounds |  |  | Site of tissue biopsies | Serological findings |  |  |  |  |  |  |  |  |  |
| --- | --- | --- | --- | --- | --- | --- | --- | --- | --- | --- | --- | --- | --- |
| No. | Age | Sex |  | SS-A<br>/Ro | SS-B<br>/La | IgG | IgG4 | IgA | IgE | IgM | CRP | RF | ANA |
| LN1 | 61 | M | Cervical lymph node | - | - | 6569 | ND | 141 | 1049 | 14 | ND | ND | + |
| LN2 | 60 | M | Cervical lymph node | - | - | 5188 | ND | 185 | ND | 40 | ND | ND | + |
| LN3 | 71 | M | Aortic lymph node | - | - | 1150 | 215 | 142 | 105 | 85 | 0.02 | 4 | 320 |
| LN4 | 61 | M | Aortic lymph node | - | - | 4970 | 773 | 120 | 706 | 54 | 0.05 | ND | ± |
| LN5 | 36 | M | Inguinal lymph node | - | - | 2837 | 963 | 406 | 1702 | 190 | 0.8 | ND | ± |
| LN6 | 67 | M | Inguinal lymph node | - | - | 2619 | 1130 | 128 | ND | 30 | 0.14 | 5 | ND |

**Table S3.**
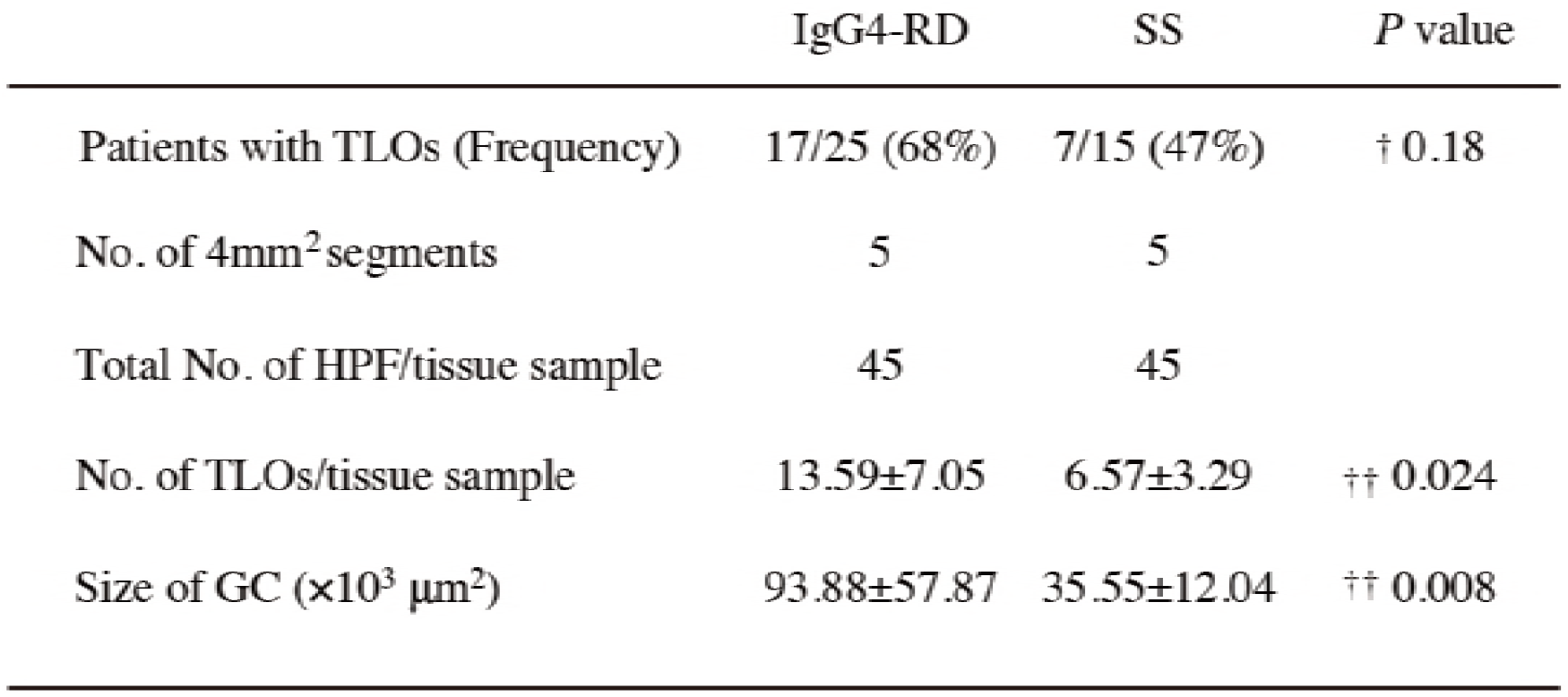
Difference in TLOs between patients with IgG4-RD and SS. **†**Significance of the difference between the two groups was determined by *X*^2^ tests. †† Significance of the difference between the two groups was determined by the Student *t* test.

|  | IgG4-RD | SS | *P* value |
| --- | --- | --- | --- |
| Patients with TLOs (Frequency) | 17/25 (68%) | 7/15 (47%) | † 0.18 |
| No. of 4mm<sup>2</sup> segments | 5 | 5 |  |
| Total No. of HPF/tissue sample | 45 | 45 |  |
| No. of TLOs/tissue sample | 13.59±7.05 | 6.57±3.29 | †† 0.024 |
| Size of GC (×10<sup>3</sup> μm<sup>2</sup>) | 93.88±57.87 | 35.55±12.04 | †† 0.008 |

